# The ciliary protein IFT88 controls post-natal cartilage thickness and influences development of osteoarthritis

**DOI:** 10.1101/2020.07.29.225599

**Authors:** CR Coveney, L Zhu, J Miotla-Zarebska, B Stott, I Parisi, V Batchelor, C Duarte, E Chang, E McSorley, TL Vincent, AKT Wann

## Abstract

Mechanical and biological cues drive cellular signalling in cartilage development, health, and disease. Proteins of the primary cilium, implicated in transduction of biophysiochemical signals, control cartilage formation during skeletal development, but their influence in post-natal cartilage remains unknown. *Ift88*^*fl/fl*^ and *AggrecanCreER*^*T2*^ mice were crossed to create a cartilage-specific, inducible knockout mouse *AggrecanCreER*^*T2*^*;Ift88*^*fl/fl*^. Tibial articular cartilage (AC) thickness was assessed, through adolescence and adulthood, by histomorphometry and integrity by OARSI score. *In situ* mechanisms were investigated by immunohistochemistry (IHC), RNA scope and qPCR of micro-dissected cartilage. OA was induced by surgical destabilisation (DMM). Mice voluntarily exercised using wheels. Deletion of IFT88 resulted in progressive reductions in medial AC thickness during adolescence, and marked atrophy in adulthood. At 34 weeks of age, medial thickness was reduced from 104.00μm, [100.30-110.50, 95% CI] in *Ift88*^*fl/fl*^ to 89.42μm [84.00-93.49, 95% CI] in *AggrecanCreER*^*T2*^*;Ift88*^*fl/fl*^ (p<0.0001), associated with reductions in calcified cartilage. Occasionally, atrophy was associated with complete, spontaneous, medial cartilage degradation. Following DMM, *AggrecanCreER*^*T2*^*;Ift88*^*fl/fl*^ mice had increased OA scores. Atrophy in mature AC was not associated with obvious increases in aggrecanase-mediated destruction or chondrocyte hypertrophy. *Ift88* expression positively correlated with *Tcf7l2*, connective tissue growth factor (*Ctgf*) and *Enpp1*. RNA scope revealed increased hedgehog (Hh) signalling (*Gli1)*, associated with reductions in *Ift88*, in *AggrecanCreER*^*T2*^*;Ift88*^*fl/fl*^ cartilage. Wheel exercise restored both AC thickness and levels of Hh signalling in *AggrecanCreER*^*T2*^*;Ift88*^*fl/fl*^. Our results demonstrate that IFT88 is chondroprotective, regulating AC thickness, potentially by thresholding a Hh response to physiological loading that controls cartilage calcification.

## Introduction

Articular cartilage absorbs and transmits mechanical loads generated by muscle contraction and weight-bearing during physical activity. Cartilage can be broadly divided into a non-calcified and a calcified zone adjacent to bone; its total thickness is allometrically scaled to the organism size ensuring chondrocytes experience similar force irrespective of animal mass (1). Articular cartilage is extremely mechanosensitive, with chondrocytes closely monitoring and remodelling their extracellular matrix in response to physiological loads (2, 3). Physiological mechanics are critical for cartilage development and homeostasis (4), their loss leading to thinning (atrophy) of cartilage (5, 6). Pathological, supraphysiological mechanical loads lead to osteoarthritis (OA) development, where degradation of the cartilage occurs with loss of integrity of the articular surface (6). The cellular response to mechanical force includes the release of fibroblast growth factor 2 (FGF2), transforming growth factor beta (TGFβ) from the matrix upon mechanical load, and activation of Hedgehog (Hh) ligand, indian hedgehog (IHH) (7-10). A number of other pathways are implicated in cellular mechanotransduction including connexin and ion channel opening and integrin activation. How chondrocytes integrate these cues as cartilage matures through adolescence, adapting to prepare for life-long loading, is far from understood.

In common with most cells of the body, articular chondrocytes express a single immotile primary cilium (11); a microtubule-based organelle reliant on intraflagellar transport (IFT) proteins, including IFT88. Components of the hedgehog (Hh) pathway localise to the cilium, supporting bi-directional modulation of signalling (12). Cilia have been directly linked with other growth factors signalling pathways such as TGFβ signalling, but also indirectly with a large, and growing, list of signalling pathways (13), many of which are pertinent to cartilage health. The primary cilium has also been coined a ‘mechanosensor’ (14). *In vitro* experiments in chondrocytes have implicated ciliary IFT88 both with compression-induced production of extracellular matrix proteins (15) and impaired clearance of aggrecanases, resulting in exacerbated degradation of aggrecan (16).

Developmental mutations in ciliary genes, including IFT88, result in impaired embryonic patterning arising from disrupted Hh signalling (17). Cartilage specific deletion of IFT88 results in disrupted long bone and articular cartilage formation (18). The cartilage-specific deletion of IFT80, in the first 2 weeks of post-natal life, resulted in thicker articular cartilage (19). However, despite its putative influence over a range of processes that regulate health and disease in cartilage, we have very limited direct knowledge about the post-developmental influence of ciliary IFT. We hypothesised that, ciliary IFT88 retains a crucial role in mediating adult cartilage homeostasis.

## Results

### Deleting IFT88 in chondrocytes results in thinner medial articular cartilage in adolescence

To confirm inducible tissue specific Cre recombination, the *AggrecanCreER*^*T2*^ line was crossed with a TdTomato reporter and tamoxifen given at 4 and 8 weeks of age. Cre recombination was activated in superficial hip articular cartilage chondrocytes (Supplementary Fig. 1A), knee articular cartilage chondrocytes and in menisci, 2 weeks later (n=3) (Fig. 1A).

**Figure 1.**
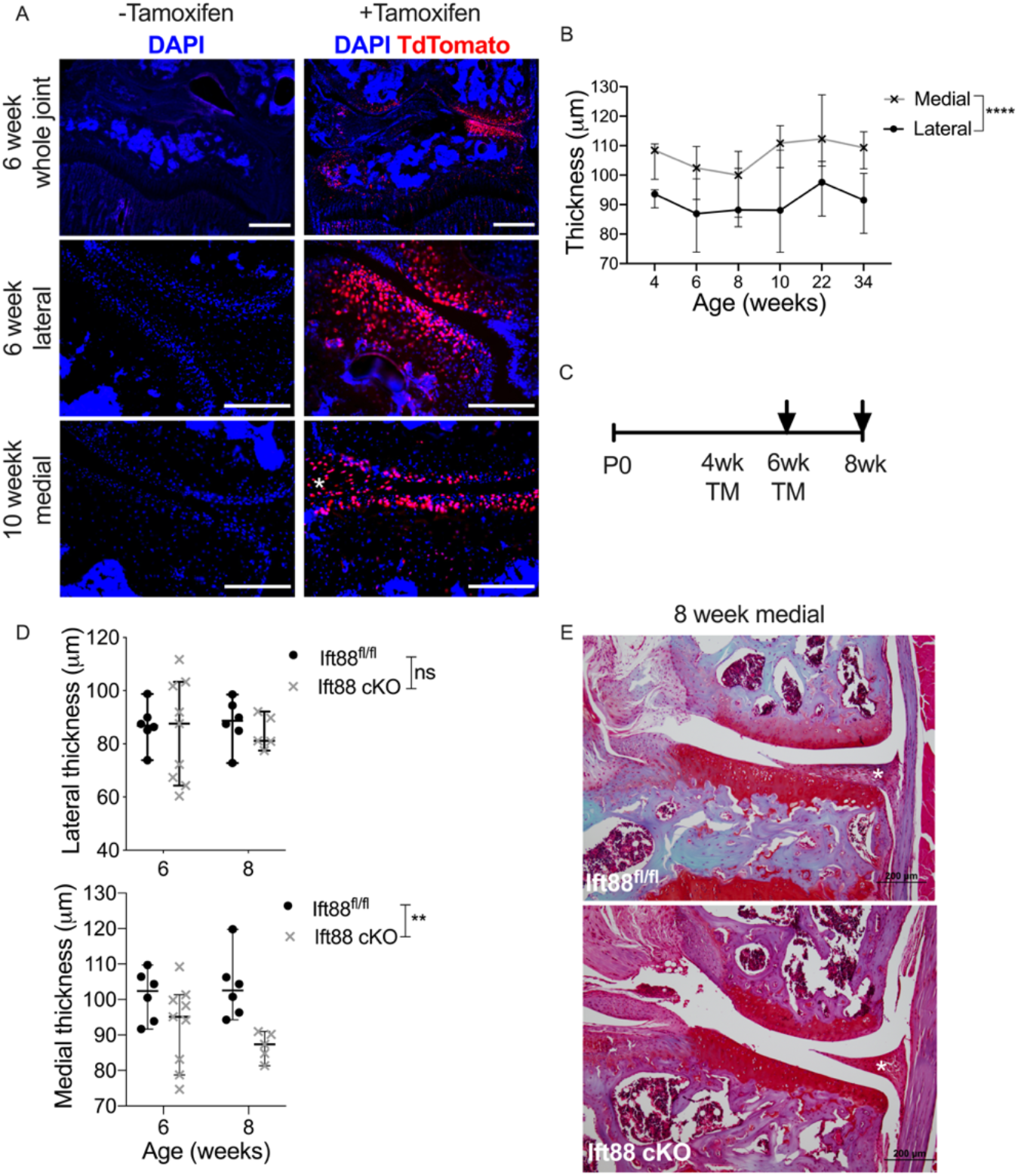
Deleting IFT88 in adolescent chondrocytes results in thinner articular cartilage. **A**, *AggrecanCreER*^*T2*^ animals were crossed with TdTomato reporter animals. Cryosections of knee joints were counterstained with nuclear stain DAPI. Animals received tamoxifen at 4 and 8 weeks of age, (scale bar= 200μm). White star labels the medial meniscus. **B**, Median lateral and medial cartilage thickness from control animals at 4, 6, 8, 10, 22 and 34 weeks of age. **C**, Experimental schematic showing age of tamoxifen administration (TM) and collection. Arrows denote collection points. **D**, Lateral and medial cartilage thickness in control (*Ift88*^*fl/fl*^) and *AggrecanCreER*^*T2*^*;Ift88*^*fl/fl*^ animals at 6 and 8 weeks of age. **E**, Safranin O stained images of representative 8-week old medial cartilage two weeks post-TM, (scale bar= 200μm). White star labels the medial meniscus. Points represent median +/- 95% confidence intervals. Analysed by two-way ANOVA, **p<0.01, ****p<0.0001.

Articular cartilage thickness on the medial tibial plateau was thicker than lateral throughout adolescence and adulthood (****p<0.0001) (Fig. 1B). Deletion of IFT88 was conducted at 4 or 6 weeks, with analysis 2 weeks later (Fig. 1C). Deletion at 6 weeks of age resulted in statistically significantly reduced median (lateral and medial combined) cartilage thickness (95.58μm, 95% CI [92.08, 99.68] in control to 82.96μm, 95% CI [81.13, 91.56] in *AggrecanCreER*^*T2*^*;Ift88*^*fl/fl*^ mice). No such effect was observed when IFT88 was deleted at 4 weeks of age (Supplementary Fig. 1B). Considering each plateau separately, deletion of IFT88 resulted in statistically significantly reduced medial cartilage thickness at 6 weeks (median thickness 102.42μm 95% CI [91.7, 109.7] in control and 94.25μm; 95% CI [78.74, 101.3] in *AggrecanCreER*^*T2*^*;Ift88*^*fl/fl*^ mice) and at 8 weeks of age (median thickness 102.57μm 95% CI [94.3, 119.8] in control, 87.36μm 95% CI [81.35, 90.97] in *AggrecanCreER*^*T2*^*;Ift88*^*fl/fl*^ mice) (Fig. 1D and E). Lateral cartilage thickness was unaffected (Fig. 1D).

### IFT88 deletion leads to thinner calcified articular cartilage

Thinner articular cartilage has been associated with acceleration of programmed hypertrophy upon post-natal activation of hedgehog (Hh) signalling (25). To assess the effects of IFT88 deletion on maturing cartilage we measured thickness of non-calcified and calcified cartilage (Fig. 2A and Supplementary Fig. S2A). In control mice, the proportion of calcified articular cartilage increases from 25% at 4 weeks of age, to 60% by 22 weeks of age (Fig. 2B). Deletion of IFT88 resulted in thinner calcified cartilage; calcified cartilage median thickness at 6 weeks of age (37.52μm 95% CI [29.71, 45.17] in control, 29.96μm 95% CI [15.46, 34.50] in *AggrecanCreER*^*T2*^*;Ift88*^*fl/fl*^) and at 8 weeks of age (48.43μm 95% CI [43.62, 54.64] in control, 41.29μm 95% CI [34.93, 44.17] in *AggrecanCreER*^*T2*^*;Ift88*^*fl/fl*^). Thickness of non-calcified cartilage was not statistically significantly different (Fig. 2C). Separation of measurements by lateral and medial plateaus revealed reductions in calcified cartilage that were observable across both plateaus and timepoints, albeit statistically significant only on the medial plateau at 8 weeks of age (Fig. 2D). Histological measurements of the subchondral bone (Supplementary Fig. 2B) revealed bone thickness approximately doubled between 6 and 8 weeks of age in control mice (Fig. 2E). However, deletion of IFT88 did not result in enhanced thickening or increased TRAP staining (Supplementary Fig. S2C), indicative of bone remodelling, suggesting very little chondro/osteoclastic activity in the calcified cartilage or subchondral bone at these timepoints.

**Figure 2.**
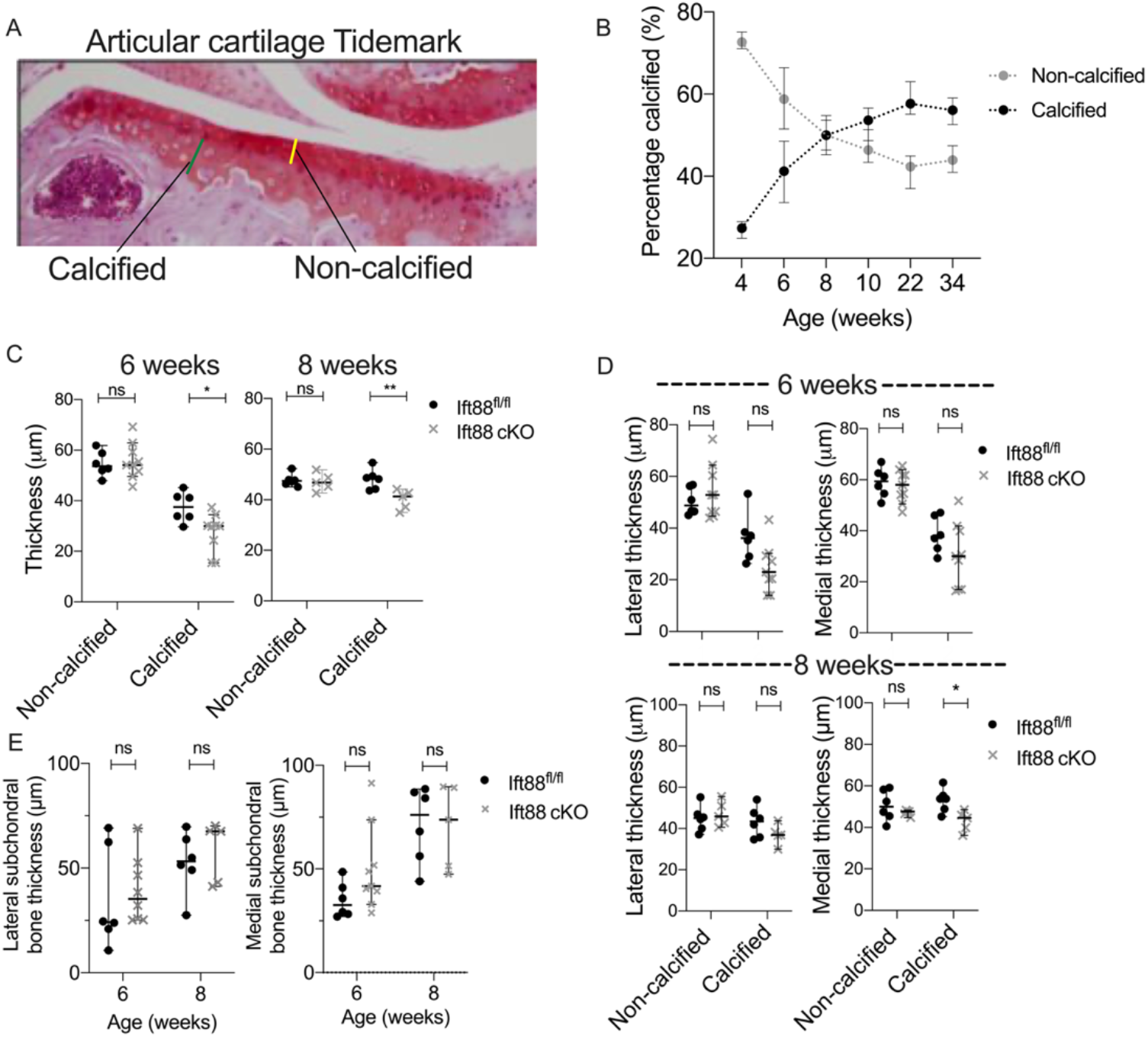
IFT88 deletion in adolescence leads to thinner calcified articular cartilage. **A**, Safranin O stained histological sections of articular cartilage showing non-calcified (yellow line) and calcified (green line) cartilage regions, demarcated by tidemark. **B**, Percentage ratios of calcified and uncalcified articular cartilage with age. At 4 weeks, n=3. In all other groups, n=6-19. **C**, Average of median medial and lateral non-calcified and calcified cartilage thickness at 6 and 8 weeks of age. **D**, Median non-calcified and calcified cartilage thickness of lateral and medial plateaus at 6 and 8 weeks of age. **E**, Median subchondral bone thickness in 6 and 8 week old joints on medial and lateral plateaus. Points represent median +/- 95% confidence intervals. Analysed by two-way ANOVA, *p<0.05, **p<0.01. Minimum n of 5 joints in any group.

### Deletion of IFT88 at 8 weeks of age results in atrophy of medial articular cartilage, associated with reduced calcified cartilage

Progressive calcification of articular cartilage continued between 8 and 22 weeks of age (Fig. 2B). Next, we administered tamoxifen at 8 weeks of age collecting joints at 10, 22 and 34 weeks of age (Fig. 3A). Deletion of IFT88 resulted in a 10-20% reduction in thickness of tibial cartilage at every time point (Fig. 3B and C). Averaging across both plateaus, thinning was not cumulative over time and occurred without obvious surface fibrillation, thus representing “atrophy” rather than “degeneration” (3, 5, 26, 27). Thinning was statistically significant at 22 and 34 weeks on the medial plateau. By 34 weeks of age, median thickness dropped from 104.00μm (95% CI [100.30, 110.50]) in control to 89.42μm (95% CI [84.00, 93.49]) in *AggrecanCreER*^*T2*^*;Ift88*^*fl/fl*^ (Fig. 3D). Atrophy was more modest on the lateral plateaus, and not apparent until 22 weeks of age (Supplementary Fig. 3A). Deletion of IFT88 did not statistically significantly affect non-calcified cartilage thickness on either plateau (Fig. 3E, left panels). In contrast, statistically significant reductions in calcified cartilage thickness were observed on both the medial (58.72μm, 95% CI [54.34, 65.05] in control to 45.62μm, 95% CI [39.32, 52.49]) in *AggrecanCreER*^*T2*^*;Ift88*^*fl/fl*^) and lateral plateaus (Fig. 3E, right panels). Cartilage thinning and accelerated hypertrophy with reduction of chondrocyte number has been observed upon hedgehog reactivation (25). However, no statistically significant reductions in cell number or density were observed in 10 week old *AggrecanCreER*^*T2*^*;Ift88*^*fl/fl*^ mice (Supplementary Fig. 3B). TUNEL staining revealed limited apoptosis in both control and *AggrecanCreER*^*T2*^*;Ift88*^*fl/fl*^ mice (Supplementary Fig. 3C).

**Figure 3.**
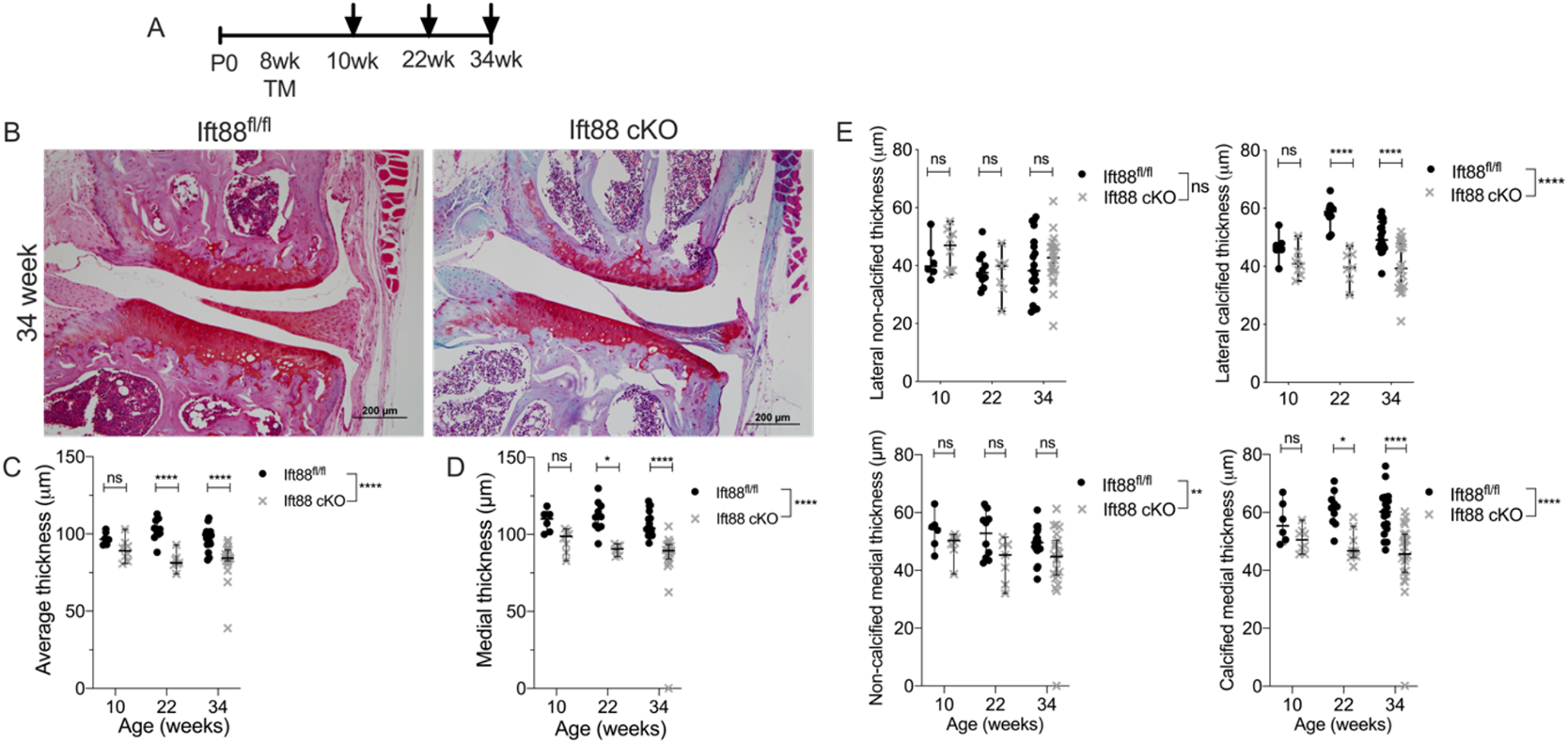
Deletion of IFT88 at 8 weeks of age results in atrophy of articular cartilage associated with reductions in calcified cartilage. **A**, Schematic representation of experimental timeline (TM= tamoxifen). Arrows denote collection points. **B**, Safranin O stained representative histological sections of medial compartment of 34-week old control and *AggrecanCreER*^*T2*^*;Ift88*^*fl/fl*^ animals (scale bar= 200μm). **C**, Average of median (both compartments) cartilage thickness measurements of mice aged 10, 22 and 34 weeks of age. **D**, Medial cartilage thickness at 10, 22 and 34 weeks of age. **E**, Lateral and medial non-calcified and calcified cartilage thickness. Points represent median +/- 95% confidence intervals. Analysed by two-way ANOVA, **p<0.01, ***p<0.001, ****p<0.0001. 10 weeks, *Ift88*^*fl/fl*^ n= 6, *AggrecanCreER*^*T2*^*;Ift88*^*fl/fl*^ n=8; 22 weeks, *Ift88*^*fl/fl*^ n= 10, *AggrecanCreER*^*T2*^*;Ift88*^*fl/fl*^ n=7; 34 weeks, *Ift88*^*fl/fl*^ n= 19, *AggrecanCreER*^*T2*^*;Ift88*^*fl/fl*^ n=22.

### Atrophy of the medial cartilage predisposes joints to spontaneous osteoarthritis

At 34 weeks of age, *AggrecanCreER*^*T2*^*;Ift88*^*fl/fl*^ mice developed spontaneous medial compartment cartilage damage associated with osteophyte formation, indicating development of osteoarthritis (Fig. 4A). OARSI scoring revealed a statistically significant increase in disease scores at 34 weeks (Fig. 4B), but none at 22 weeks of age (Supplementary Fig. 4A). The highest OARSI scores were found on the medial plateau (Fig. 4C). There was no difference in osteophyte scores except in the presence of severe cartilage damage (Fig. 4D and Supplementary Fig. 4B). There were no changes in subchondral bone thickness with IFT88 deletion at either 10 or 22 weeks of age. By 34 weeks of age, thinning of cartilage was associated with loss of subchondral bone in *AggrecanCreER*^*T2*^*;Ift88*^*fl/fl*^ animals (Fig. 4E and Supplementary Fig. 4C). Trabecular bone density (BV/TV) in the epiphysis was not affected by deletion of IFT88 (Supplementary Fig. 4D).

**Figure 4.**
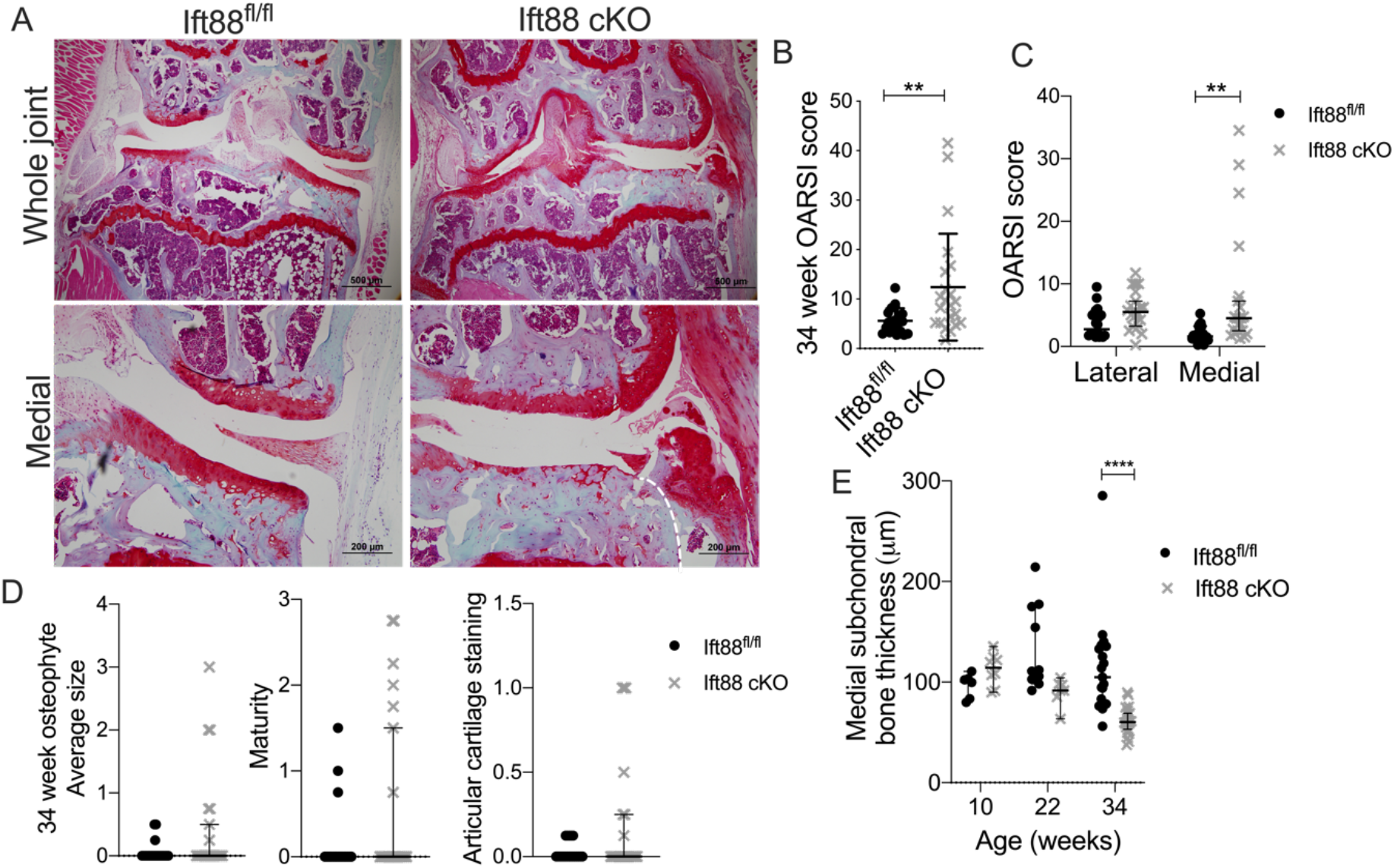
Spontaneous osteoarthritis in IFT88 *AggrecanCreER*^*T2*^*;Ift88*^*fl/fl*^ at 34 weeks of age. **A**, Safranin O stained histological sections of whole joint (scale bar= 500μm) and medial compartment (scale bar= 200μm) of 34-week old control and *AggrecanCreER*^*T2*^*;Ift88*^*fl/fl*^ animals, selecting an example displaying medial OA rather than representative shown in Figure 3. White dashed line demarcates formation of osteophyte. **B**, Summed modified OARSI scores in sections from 34-week old animals. Analysed by Mann-Whitney test, **p<0.01. **C**, Summed modified OARSI scores from lateral and medial plateaus in sections from 34-week old animals. **D**, Osteophyte size, osteophyte maturity, and staining of articular cartilage scores at 34 weeks of age. **E**, Medial subchondral bone thickness. 34 weeks: *Ift88*^*fl/fl*^ n= 19, *AggrecanCreER*^*T2*^*;Ift88*^*fl/fl*^ n=22. Points represent median +/- 95% confidence intervals. Analysed by two-way ANOVA unless stated otherwise, ****p<0.0001.

### Deletion of IFT88 exacerbates osteoarthritis severity after surgical joint destabilisation

Mice were given tamoxifen at 8 weeks of age, and surgical destabilisation of the medial meniscus (DMM) was performed at 10 weeks. At 8 or 12 weeks post-surgery, animals were culled and joints collected, along with respective sham operated mice (Fig. 5A). Coronal histological sections of joints were assessed using summed OARSI scoring (Fig. 5B). Deletion of IFT88 resulted in exacerbated disease at 12 weeks post-DMM (*Ift88*^*fl/fl*^ 22.08± 9.30, *AggrecanCreER*^*T2*^*;Ift88*^*fl/fl*^ 29.83± 7.69, p<0.05, n=15 for both groups) (Fig. 5C). No differences in osteophyte size (Fig. 5D and Supplementary Fig. 5A), maturity or staining (Supplementary Fig. 5B-E) were observed between control and *AggrecanCreER*^*T2*^*;Ift88*^*fl/fl*^ animals at either time-point post-DMM. Further, no difference in BV/TV between control and *AggrecanCreER*^*T2*^*;Ift88*^*fl/fl*^ epiphyses 12 weeks following DMM was observed (Supplementary Fig. 5F).

**Figure 5.**
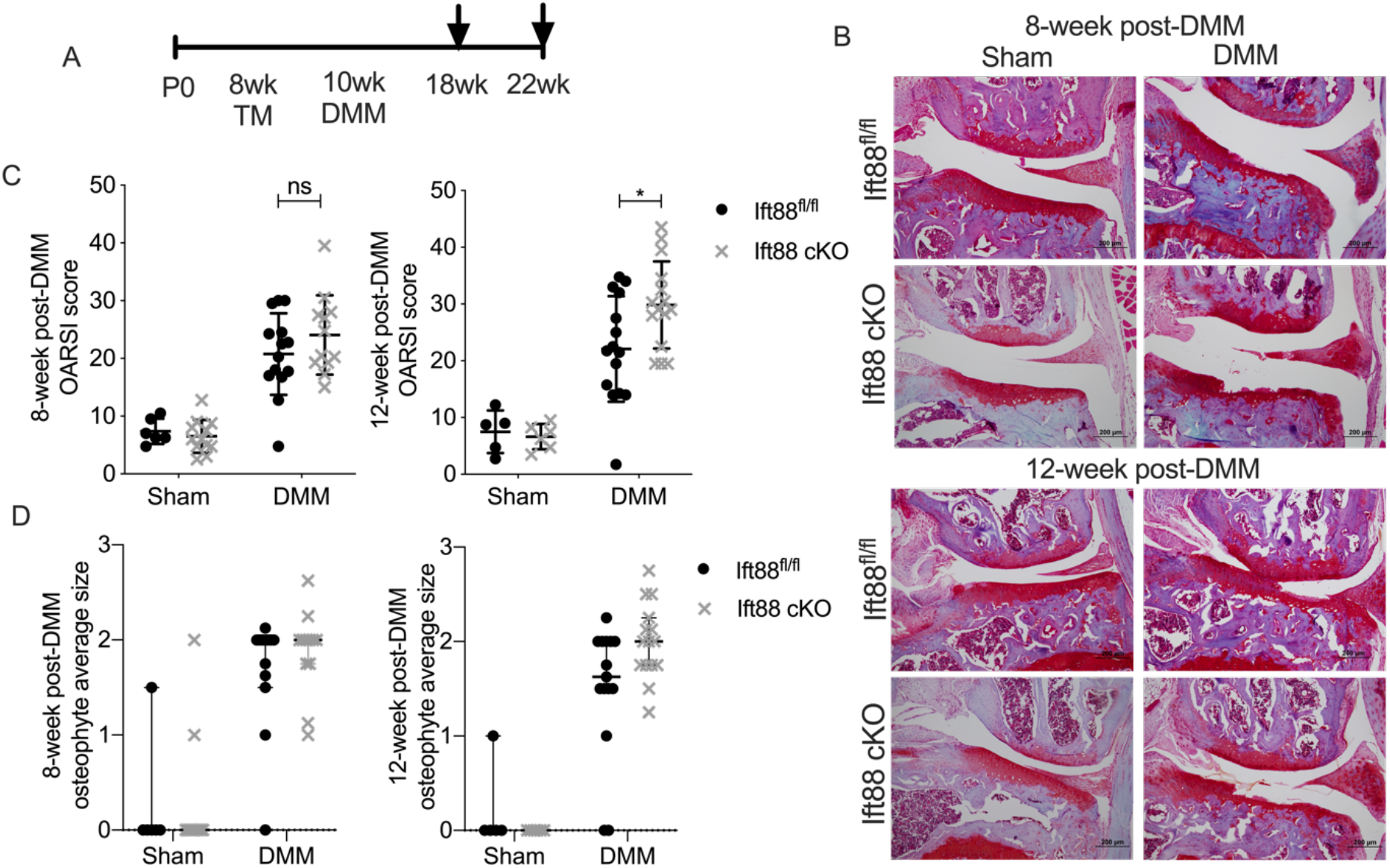
Deletion of IFT88 results in exacerbated disease 12 weeks after surgical destabilisation. **A**, Schematic representation of experimental timeline for DMM surgery (TM= tamoxifen). Arrows denote collection points. **B**, Safranin O stained histological sections of medial compartments from sham and DMM animals operated animals 8 and 12 weeks post DMM surgery. Scale bars= 200μm. **C**, Summed modified OARSI scores in sections from sham and DMM operated knees 8 and 12 weeks post-DMM. Minimum sham surgeries in any group n= 5. **D**, Osteophyte average size 8 and 12 weeks post-DMM. 8 weeks post-DMM; *Ift88*^*fl/fl*^ n= 14, *AggrecanCreER*^*T2*^*;Ift88*^*fl/fl*^ n=12. 12 weeks post-DMM; *Ift88*^*fl/fl*^ n= 15, *AggrecanCreER*^*T2*^*;Ift88*^*fl/fl*^ = 15. Points represent median +/- 95% confidence intervals. Analysed by two-way ANOVA, *p<0.05.

### Articular cartilage atrophy is not associated with markers of matrix catabolism, or hypertrophy

To assess if IFT88 deletion increased catabolism of aggrecan, control and *AggrecanCreER*^*T2*^*;Ift88*^*fl/fl*^ joints were probed for protease-generated, neoepitopes as described previously (22), using an antibody raised against the synthetic epitope NITEGE, thresholded to IgG control (Supplementary Fig. 6A, right-hand panel). NITEGE signal (Fig. 6A) was weak with respect to positive control sections (Supplementary Fig. 6A left-hand panel) and no differences were observed when comparing controls with *AggrecanCreER*^*T2*^*;Ift88*^*fl/fl*^. No differences in type X collagen (ColX) expression, a marker of Hh-driven chondrocyte hypertrophy (28), were observed by immunohistochemistry (Fig. 6B, IgG control Supplementary Fig. 6B).

**Figure 6.**
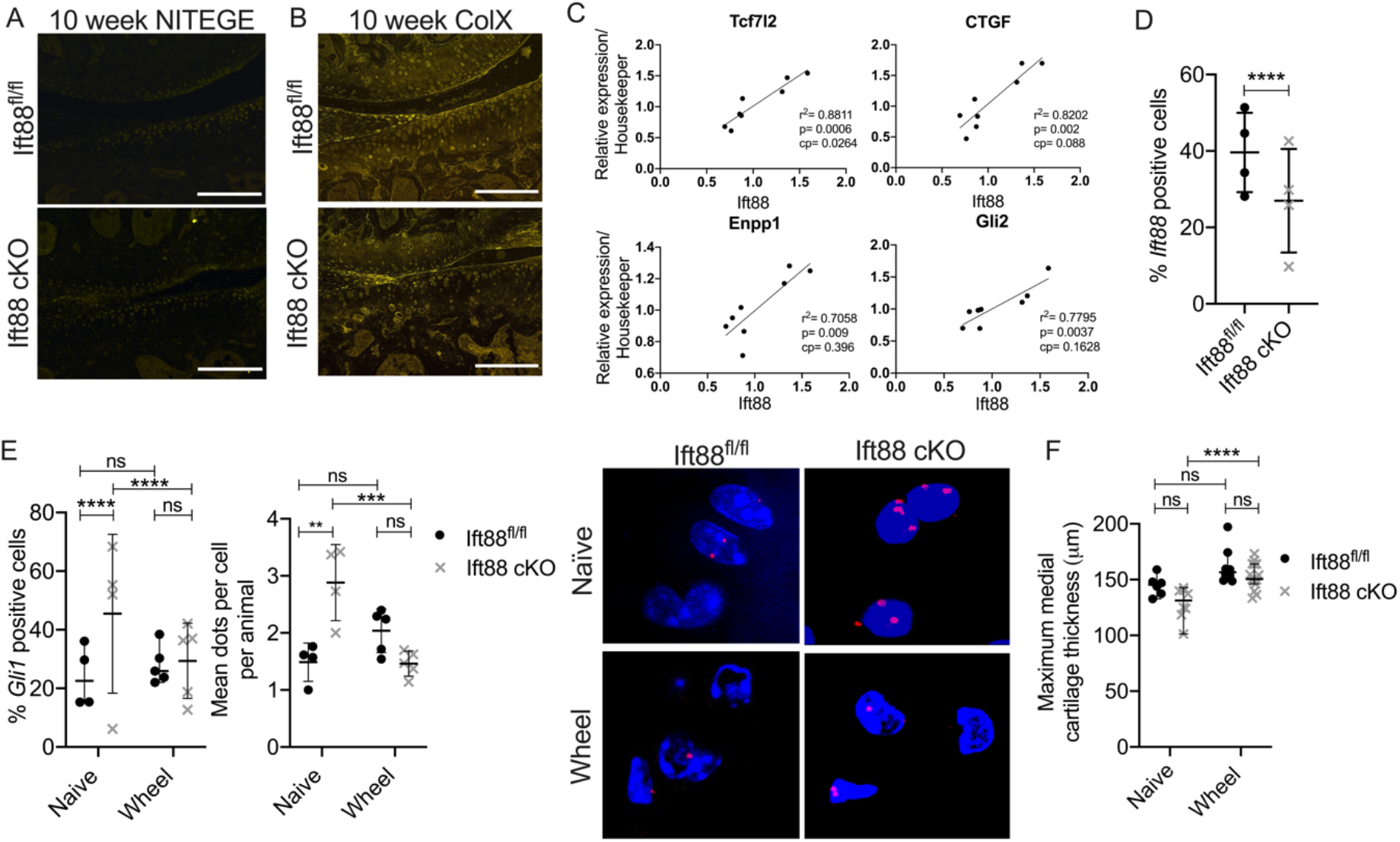
Wheel exercise rescues atrophy and supresses increased chondrocyte hedgehog signalling. **A**, Representative immunofluorescent staining of aggrecan neoepitope NITEGE (n=6 in all groups). Scale bars=200μm. **B**, Representative immunofluorescent staining of type X collagen (n=6 in all groups). Scale bars=200μm. **C**, RNA extracted from micro-dissected articular cartilage was analysed by qPCR to identify genes correlating with *Ift88* expression following normalisation to *Gapdh* and *Hprt*. Linear regression was performed and statistical significance assessed following Bonferroni correction (cp= corrected p value). *Tcf7l2* positively correlated with *Ift88* expression before and after correction (p=0.0006, cp=0.0264), whilst *Ctgf, Enpp1*, and *Gli2* correlated positively before correction (Supplementary Table 1). **D**, RNAscope analysis of articular cartilage to assess *Ift88* expression. (Analysed by Fisher’s exact test, ****p<0.0001), *Ift88*^*fl/fl*^ n=4, *AggrecanCreER*^*T2*^*;Ift88*^*fl/fl*^ n=4. **E**, RNAscope analysis of articular cartilage to assess *Gli1* expression. (Analysed by Fisher’s exact test, ****p<0.0001, Supplementary Fig. 6E), and the mean dots per cell per animal. *Ift88*^*fl/fl*^ naïve: n=4, wheel: n=5, *AggrecanCreER*^*T2*^*;Ift88*^*fl/fl*^ naïve: n=4, wheel: n=5. Representative images of *Gli1* positive nuclei. **F**, Maximum medial cartilage thickness in naïve (non-exercised) and wheel exercised mice. *Ift88*^*fl/fl*^ n=10, *AggrecanCreER*^*T2*^*;Ift88*^*fl/fl*^ n=15. Points represent median +/- 95% confidence intervals. Analysed by two-way ANOVA, *p<0.05, **p<0.01, ***p<0.001.

### Bulk RNA analysis implicates *Tcf7l2* in cartilage atrophy

RNA was isolated from micro-dissected articular cartilage (Supplementary Fig. 6C) from 10 week old control and *AggrecanCreER*^*T2*^*;Ift88*^*fl/fl*^ animals, 2 weeks after tamoxifen treatment. 44 candidate genes, associated with chondrocyte biology and/or implicated in primary ciliary signalling, were measured by qPCR. Due to the mosaic Cre activity (Fig. 1A) and sample pooling we conducted a correlative analysis with *Ift88* expression, normalised to the average of the two most stable housekeeper genes, *Hprt* and *Gapdh* (See Supplementary Table 1 for all genes). Expression of the majority of candidate genes did not correlate with *Ift88*, including *Adamts5* and *ColX* (Supplementary Fig. 6D) in line with our immunohistochemistry (Fig. 6A and B). *Tcf7l2 (29)* was the only gene found to be statistically significantly correlated with *Ift88* expression after Bonferroni correction (p=0.0006, cp=0.026, r^2^=0.8811). *Ctgf, Tgfbr3, Gli2* and the regulator of cartilage calcification *Enpp1* (30) positively correlated with *Ift88* expression before correction (p=0.002, p=0.0037, and p=0.009 respectively) (Fig. 6C and Supplementary Fig. 6C), but not after correction. Expression of *Gli1* and *Ptch1*, indicators of Hh pathway activation, were not correlated with *Ift88* expression (Supplementary Table 1).

### RNA scope analysis reveals reduction in IFT88 expression in *AggrecanCreER*^*T2*^*;Ift88*^*fl/fl*^ cartilage

To overcome the limitations of bulk RNA analyses, RNAScope analysis was performed on cryosections of 10 week old tibial articular cartilage from 4 control and 4 *AggrecanCreER*^*T2*^*;Ift88*^*fl/fl*^ mice to assess *Ift88* expression, *in situ*. RNA scope revealed 45.18% of cells were *Ift88* positive in control articular cartilage in comparison with 27.78% in *AggrecanCreER*^*T2*^*;Ift88*^*fl/fl*^ indicating a 40% reduction in *Ift88* positive cells (p<0.0001, Fischer’s exact test) (Fig. 6D, images and contingency data for statistical comparisons shown in Supplementary Fig. 6E).

### Cartilage atrophy and associated increases in hedgehog signalling *in situ* are rescued by wheel exercise

RNAScope was performed to investigate *Gli1* expression, a marker of Hh pathway activity in the same sections where IFT88 was reduced. 23.63% of nuclei were *Gli1* positive in control articular cartilage in comparison with 50.42% in *AggrecanCreER*^*T2*^*;Ift88*^*fl/fl*^ (Fig. 6E). This increase in *Gli1* positive cell count, indicative of increased Hh pathway activity, was statistically significant (p<0.0001, n=4 in both groups, raw counts shown in Supplementary Fig. 6E). Analysing *Gli1* in terms of signal per cell also revealed statistically significantly (p<0.01, n=4 for both groups) higher expression of *Gli1* in *AggrecanCreER*^*T2*^*;Ift88*^*fl/fl*^ (2.88) compared with controls (1.49), as shown in representative images (Fig. 6E). Mice were given free access to wheel exercise for 2 weeks immediately following tamoxifen injection at 8 weeks of age (over the same time period atrophy develops). In these experiments, no measurable difference in articular cartilage thickness was observed comparing control mice with *AggrecanCreER*^*T2*^*;Ift88*^*fl/fl*^ (Fig. 6F and Supplementary Fig. 6F). Wheel exercise did not affect the number of *Gli1* positive cells or *Gli1* expression per cell in control mice, but reduced *Gli1* expression, by both analyses, in *AggrecanCreER*^*T2*^*;Ift88*^*fl/fl*^ effectively restoring hedgehog signalling and cartilage thickness to control levels.

## Discussion

In this study we explored the influence of ciliary IFT88 in post-natal articular cartilage, *in vivo*, by depleting its expression in chondrocytes at different stages of post-natal skeletal maturation. We report rapid cartilage thinning when *Ift88* was deleted at 4, 6, and 8 weeks of age which was largely in the calcified cartilage within the medial joint compartment. Thinning was deemed to be “atrophy” rather than “degeneration” as there was no disruption to the articular surface. However, with age, cartilage atrophy was associated with increased spontaneous and DMM-induced osteoarthritis.

We speculate that the IFT88-dependent effects in the thicker medial compartment of the joint are mechanically driven (6), in a manner analogous to the bone ‘mechanostat’ proposed by Frost in 1987 (31). In essence this model ensures that the chondrocytes mechanoadapt their extracellular matrix so as to experience force within a narrow window (1). As cartilage thickness can be recovered in *Ift88* KO mice in response to wheel exercise, this suggests that ciliary IFT88 may influence, but is not solely responsible for, cartilage mechanoadaptation *in vivo*. Similar modes of action have been proposed in the context of epithelial response to renal flow (14, 32). Our study, contrasts with the observation that deletion of IFT88 during development leads to increased cartilage thickness [17], possibly indicating a changing influence for *Ift88* with skeletal maturation.

As atrophy in *Ift88* KO mice is largely restricted to the calcified cartilage, we speculate that this represents a failure of cartilage hypertrophy during maturation, in a load dependent manner. Cartilage thinning was not associated with increased matrix catabolism, enhanced subchondral bone thickness, chondro/osteoclast activity or changes to the density (BV/TV) of the epiphysis, but we can’t exclude the possibility that calcified cartilage is transitioning to bone in these mice. Calcification of cartilage in the mouse has recently been linked to ENPP1, a pyrophosphatase proposed to inhibit calcification through Hh signalling (30). Our molecular analyses indicated ENPP1 positively correlates with *Ift88* expression, implying a reduction in the levels of an inhibitor of calcification in *Ift88* KO mice. This could result in accelerated ossification analogous to that seen in the growth plates of ciliary mutants and in the articular cartilage upon post-natal activation of Hh (25).

Studies prior to this, describing congenital mutations (33) or constitutive deletions of IFT88 (17, 18, 34), demonstrate a role for IFT88 in limb and joint development. Mouse models targeting other ciliary components, KIF3A, BBS proteins, and IFT80, also implicate the ciliary machinery in musculoskeletal development (19, 28), which are phenocopied by human skeletal ciliopathies (35, 36). The most important molecular pathway associated with ciliopathy is Hh although other pathways have also been described (10, 37-41). Hh signalling largely switches off in adulthood (42), but is reactivated in OA (8). We investigated the molecular basis of IFT88-dependent cartilage atrophy by correlating expression of *Ift88* (reflecting efficiency of deletion) with 44 molecules previously associated with ciliary signalling and cartilage biology. In addition we explored Hh signalling directly by visualising *Gli1* expression in murine cartilage by RNAscope The most strongly correlated gene was transcription factor *Tcf7l2*, previously shown to influence and interact with Hh and *β*-catenin signalling pathways in cartilage (37). Other genes that correlated with Ift88, albeit significant only before correction, included *Gli2, Ctgf and Enpp1*.

Whilst gene expression analysis of micro-dissected cartilage did not show correlation of *Ift88* with classical Hh pathway molecules (*Gli1, Ptch1*), individual cell analysis by RNA scope did reveal increased *Gli1* expression in *AggrecanCreER*^*T2*^*;Ift88*^*fl/fl*^ *chondrocytes*, suggesting a reciprocal relationship between Hh signalling and *Ift88*. Therefore, we propose that loss of IFT88 disrupts ciliary-mediated repression of Hh signalling, resulting in net increases in *Gli1* expression. This is in agreement with previous studies in constitutive *Ift88* deletion (18) and also with that observed in endochondromas (43). Rescue of cartilage atrophy and normalisation of basal *Gli1* expression with voluntary wheel exercise in *AggrecanCreER*^*T2*^*;Ift88*^*fl/fl*^ mice provides critical evidence for a link to mechanical load in this model. Our data imply that in post-natal articular cartilage, ciliary IFT88 safeguards the progressive mechanoadaptation of adolescent cartilage, supporting the creation and maturation of fit-for-purpose adult articular cartilage, by ensuring appropriate levels of hedgehog signalling.

We recognise a number of limitations in our study. We induced deletion of IFT88 in chondrocytes, by expressing Cre recombinase on the aggrecan promotor to ensure good expression throughout adulthood (44). Despite this, we observed only 40% reduction in *Ift88* positive cells in the tibial articular cartilage of *AggrecanCreER*^*T2*^*;Ift88*^*fl/fl*^. The tomato reporter indicated that deletion took place in only a proportion of chondrocytes but this activity was not biased by knee compartment so is unlikely to account for the differences observed in the medial and lateral sides of the joint. We also recognise that we have not yet been able to address the question of whether primary cilia have been perturbed as a result of IFT88 deletion, due to the challenges associated with imaging cilia in cartilage, thus cannot conclude if molecular changes are direct consequence of cilia loss. We have not yet conducted deletion of *Ift88* later than 8 weeks of age to test its influence later in adulthood or assessed behaviour in *AggrecanCreER*^*T2*^*;Ift88*^*fl/fl*^ mice.

In summary, these data demonstrate that IFT88 remains highly influential in adolescent and adult articular cartilage, as a positive regulator of cartilage thickness, guiding cartilage calcification during maturation, and serving as a guardian of physiological Hh signalling in adult cartilage in response to mechanical cues.

## Materials and Methods

### Animals

*Ift88*^*fl/fl*^ mice were obtained from Jackson labs (Stock No. 022409) and maintained as the control line, and in parallel offspring were crossed with the *AggrecanCreER*^*T2*^ mouse line, *AggrecanCreER*^*T2*^*;Ift88*^*fl/fl*^ (Ift88 cKO), originally generated at the Kennedy Institute of Rheumatology (20). The TdTomato reporter mouse line *B6*.*Cg-Gt(ROSA)26Sor*^*tm14(CAG-TdTomato)Hze*^*/J* was originally from Jackson Laboratories (Stock No. 007914). For all experiments, apart from DMM and wheel exercised (male only), both sexes were used. No effect of sex was observed in data.

### Tamoxifen treatment

Tamoxifen (Sigma-Aldrich, T5648) was dissolved by sonication in 90% sunflower oil and 10% ethanol at a concentration of 20mg/ml. Tamoxifen was administered to *Ift88*^*fl/fl*^, *AggrecanCreER*^*T2*^*;Ift88*^*fl/fl*^, and *AggrecanCreER*^*T2*^; *TdTomato* mice via intraperitoneal injection on three consecutive days at 50-100mg/kg (dependent on animal weight), at either 4, 6, or 8 weeks of age dependent on experiment (n=5-22).

### Destabilisation of the medial meniscus (DMM)

Mice were given tamoxifen 2 weeks prior to surgery. Male *Ift88*^*fl/fl*^ *(control)* and *AggrecanCreER*^*T2*^*;Ift88*^*fl/fl*^ mice underwent DMM surgery at 10 weeks of age as previously described (21) or capsulotomy as Sham surgery. DMM: Mice were culled 8 (*Ift88*^*fl/fl*^ n=14, *AggrecanCreER*^*T2*^*;Ift88*^*fl/fl*^ n=12) or 12 weeks (*Ift88*^*fl/fl*^ n=15, *AggrecanCreER*^*T2*^*;Ift88*^*fl/fl*^ =15) post-DMM. Mice were anaesthetised as described previously (22).

### Histology

Knee joints were harvested into 10% neutral buffered formalin (CellPath, Newtown, UK). Joints were decalcified (EDTA), paraffin embedded, coronally sectioned through the entire depth of the joint. Sections (4μm), at 80μm intervals were stained with Safranin O.

### OARSI scoring

Safranin O-stained joint sections were assessed by a cartilage scoring system, as previously described (21), by 2, blinded, observers. Summed score method (sum of 3 highest scores per section, per joint, with a minimum of 9 sections per joint) was used.

### Osteophyte quantification

Osteophyte size, maturity, and loss of medial tibial cartilage proteoglycan staining was assessed on a 0-3 scale as described previously (23).

### Cartilage thickness measurements

Thickness measurements were taken from the average of three measurements from both the medial and lateral tibial plateaus (6 measurements), from three consecutive sections from the middle of the joint (18 measurements). The same method was used to measure non-calcified cartilage thickness in the same location as the total cartilage thickness measurement. Calcified cartilage thickness was calculated by subtracting, non-calcified cartilage from total cartilage thickness. All images were independently, double-scored; measurements performed using ImageJ (Supplementary Fig. 2C).

### Subchondral bone measurements

Subchondral bone measurements were taken as previously described (24). Briefly, using ImageJ, five measurements were taken perpendicular to the articular cartilage plateau, from the chondro-osseous junction to the top of bone marrow or to the growth plate if there was no bone marrow. Thickness was measured from at least 3 sections of each plateau of each knee.

### MicroCT BV/TV

Knee joints scanned by SkyScan1172 X-ray microtomograph in 70% ethanol (10μm/pixel). 3D parameter analysis including tissue volume, bone volume, tissue surface, bone surface, trabecular thickness, separation, and number, and trabecular pattern factor conducted using CTan (Brucker).

### Immunohistochemistry

Fixed, decalcified, unstained coronal knee sections were deparaffinised, rehydrated, quenched in 0.3M glycine and treated with proteinase K for 10 minutes. Samples underwent chondroitinase (0.1U) treatment for 30mins at 37°C blocked in 5% goat serum and 10% bovine serum albumin (BSA) in phosphate-buffered saline, and were permeabilised by 0.2% Triton X-100 for 15mins. Samples were incubated with primary antibody (anti-ColX, 1:50, Abcam, ab58632, anti-NITEGE, 1:50, Thermo Scientific, PA1-1746, IgG (1:50) control, or no primary) overnight at 4°C. Sections were washed and incubated with Alexa-conjugated 555 secondary antibodies for 30 mins. Samples were incubated with nuclear stain DAPI (1:5000), before mounting in Prolong Gold.

### RNA extraction

Articular cartilage was micro-dissected using a scalpel from the femoral and tibial plateaus (each data point, n, from two animals), and harvested directly into RNA*later* (Invitrogen). Samples were transferred to buffer RLT (Qiagen) and diced before pulverisation using a PowerGen 125 Polytron instrument (Fisher Scientific) (3×20 seconds). RNA of knee cartilage was isolated using the RNeasy Micro kit (Qiagen).

### qPCR

Using RNA isolated from micro-dissected knee cartilage, cDNA was synthesised (Applied Biosystems) and real-time quantitative PCR was performed using a Viia 7 Real-Time PCR System on 384 custom-made TaqMan microfluidic cards (ThermoFisher, 4342253). CTs were normalised to the average CT values of *Gapdh* and *Hprt*. Linear regressions were used to correlate *Ift88* expression and subsequent raw p values and corrected (Bonferroni) calculated using GraphPad Prism 9.

### RNAScope

Knee joints were harvested into ice cold 4% PFA and incubated in the fridge for 24 hours before transfer to ice cold 10%, 20% and 30% sucrose, each for 24 hours. RNAScope probes Mm-Ift88-C1 (420211-C1) and Mm-Gli1-C2 (311001-C2) were used in combination with Opal™ 570 FP14880001KT and 690 Reagent Pack FP1497001KT signal normalised to positive/negative controls. For further details on slide preparation and the RNAScope® Multiplex Fluorescent Reagent Kit v2 Assay reagents (323100) protocol, imaging, and normalisation see supplementary materials.

## Supplementary Methods

### Animals

All mice were housed in the biomedical services unit (BSU) at 4–7 per standard, individually-ventilated cages and maintained under 12-h light/12-h dark conditions at an ambient temperature of 21°C.

### RNAScope

Following sucrose gradients, Knee joints were embedded into Super Cryo Embedding Medium (C-EM001, Section-lab Co. Ltd). 8μm sections were collected using a pre-cooled (−16 °C) cryotome with Cryofilm type 3C (16UF) 2.5 cm C-FUF304. Slides were washed with PBS for 5mins, baked at 60°C for 30mins and fixed using ice cold 4% PFA for 15 mins at 4 °C. Increasing concentrations of ethanol was applied, 50%, 70%, 100%, and 100% fresh ethanol, 5mins for each gradient.

The sample was air dried for 5mins and incubated with hydrogen peroxide for 10mins. Slides were submerged twice in milli-Q water and then transferred into pre-warmed Target Retrieval Buffer (1X) (322000) in the steamer for 10mins at 75°C. Slides were washed briefly in milli-Q water before being submerged briefly in 100% ethanol and air dried for 5mins. Samples were covered in Protease III (PN 322381) and incubated at 40°C for 30mins. Slides were submerged briefly in milli-Q water.

RNAscope® Multiplex Fluorescent Reagent Kit v2 Assay reagents (323100) was subsequently followed. The whole tibial plateau was imaged using a 60x lens, 110nm/px, 381.84um x 866.71um, 5um z-stack, using a Nikon A1R HD25 or Zeiss 980 confocal microscope. Following normalisation with positive (RNAscope® 3-plex Positive Control Probe-Mm, PPIB gene, 320881) and negative (RNAscope® 3-plex Negative Control Probe-Mm, bacterial dapB gene, 320871) control probes, the number of *Ift88* and *Gli1* positive and negative nuclei, and the number of dots per cell were counted.

### Cell counting

Cartilage plateau area and cell count was conducted in ImageJ. Three sections in the middle of the joint were assessed per plateau, therefore, 6 measurements taken per joint. The assessor was blinded to genotype.

### TUNEL

*In situ* detection of apoptosis was conducted using TACS® 2 Tdt-Fluor *In Situ* apoptosis kit (Trevigen, 4812-30-K), after deparaffinising sections.

## Supplementary Figures

**S1.**
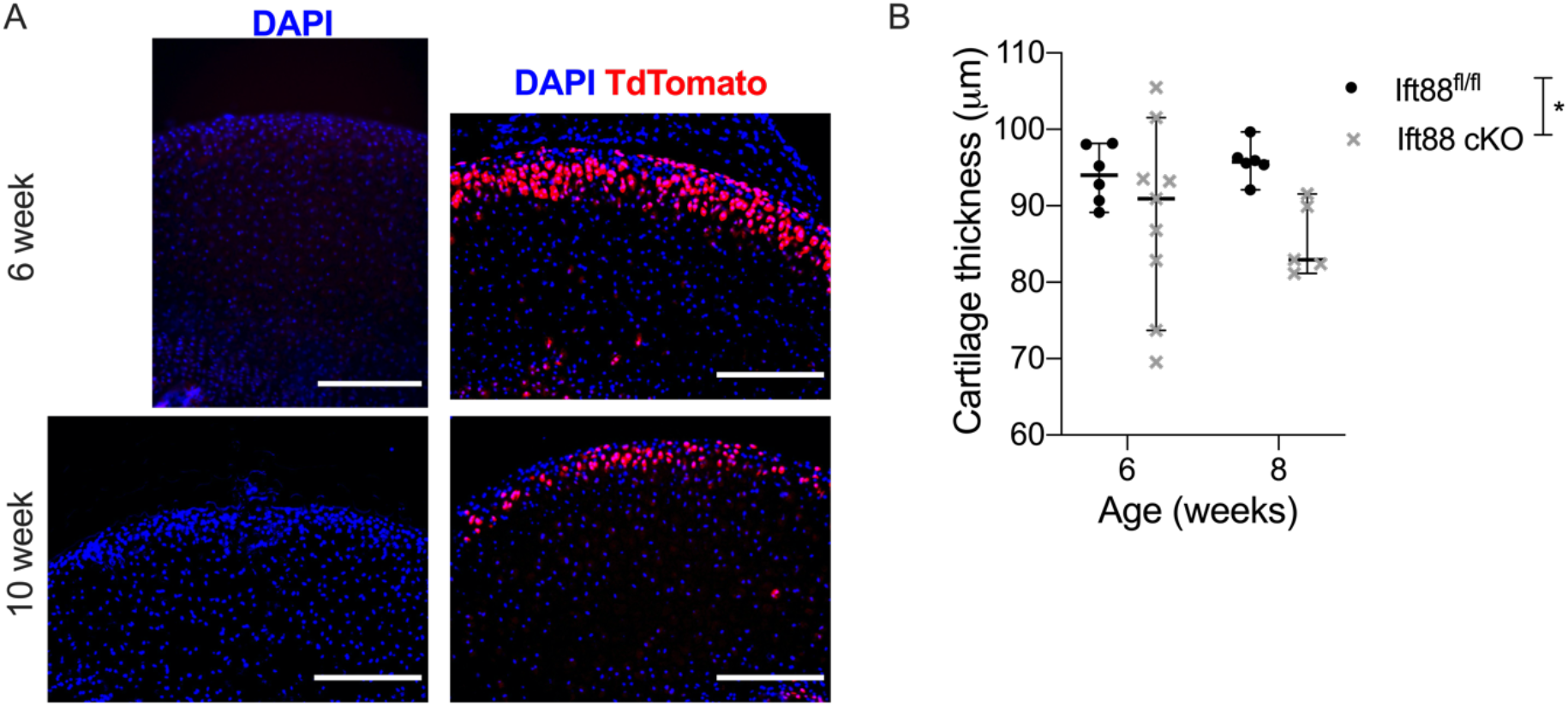
**A**, TdTomato activated by Cre recombination in 6 and 10 week old hip articular cartilage, following tamoxifen administration at 4 and 8 weeks of age respectively (scale bar=200 μm). **B**, Average cartilage thickness of *Ift88*^*fl/fl*^ and *AggrecanCreER*^*T2*^*;Ift88*^*fl/fl*^ animals (*p<0.05), two weeks post tamoxifen administered at 4 and 6 weeks of age. Points represent median +/- 95% confidence intervals. Analysed by two-way ANOVA, *p<0.05.

**S2.**
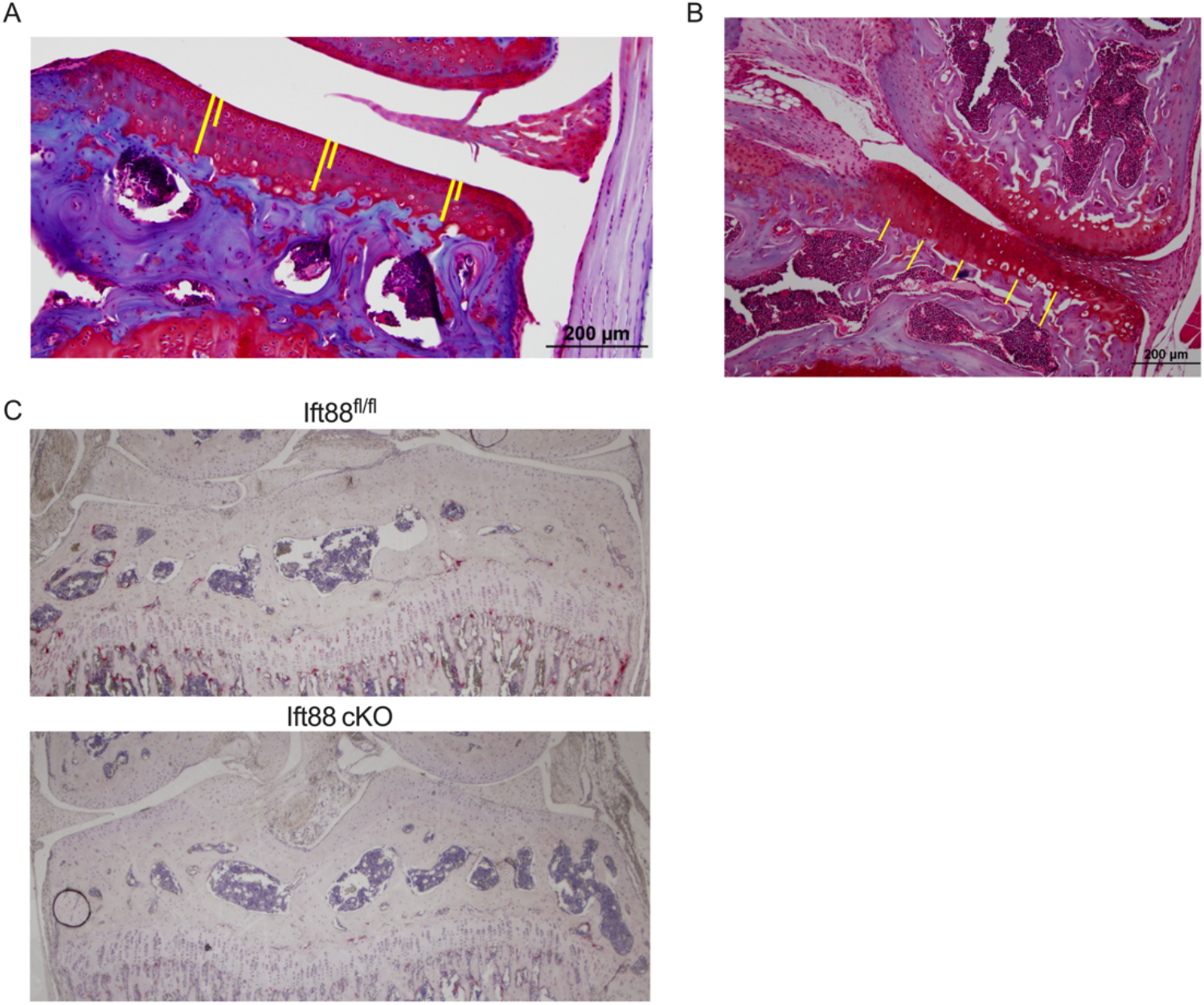
**A**, Yellow lines depict measurements taken from the total thickness of the articular cartilage and the non-calcified region demarcated by the tidemark at three points across each of the medial and lateral plateaus and averaged over three consecutive sections from the middle of the joint. **B**, Five subchondral bone measurements are taken across the plateau from the chondro-osseous junction to the top of the bone marrow and averaged across three sections from the middle of the joint (n=5-10 in any group). **C**, Osteoclastic activity in the articular cartilage was assessed by TRAP staining in control and *AggrecanCreER*^*T2*^*;Ift88*^*fl/fl*^ 8 week old mice (n=5 in both groups).

**S3.**
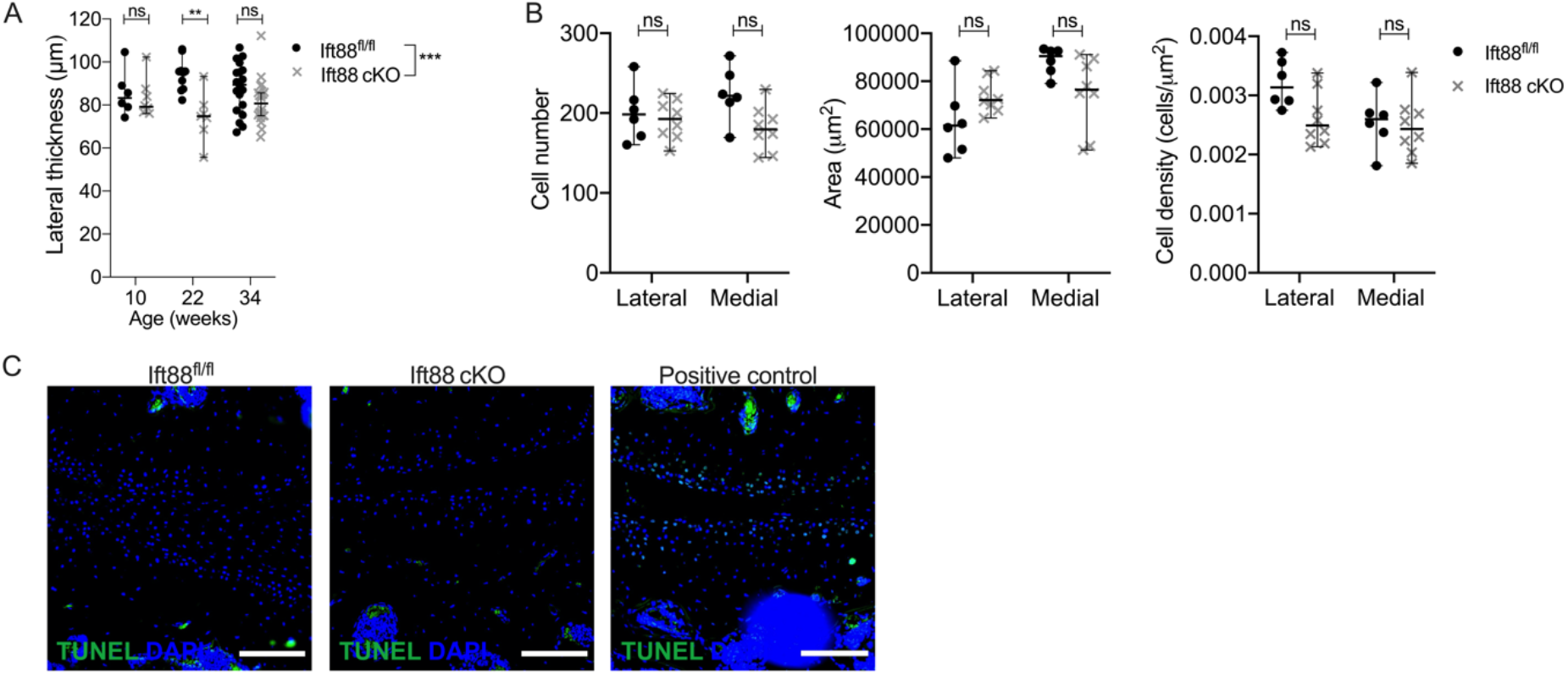
**A**, Lateral cartilage thickness 10, 22 and 34 weeks of age. **B**, Quantification of chondrocyte cell number, area, and cell density in the articular cartilage of control and *AggrecanCreER*^*T2*^*;Ift88*^*fl/fl*^ 10 week old mice, having received tamoxifen at 8 weeks of age. **C**, TUNEL (green) staining was used to assess apoptosis in control and *AggrecanCreER*^*T2*^*;Ift88*^*fl/fl*^ 10 week old mice having received tamoxifen at 8 weeks of age. Slides were counterstained with DAPI, and positive control shown (n=3 in both groups). Points represent median +/- 95% confidence intervals. Analysed by two-way ANOVA, **p<0.01, ***p<0.001. 10 weeks, *Ift88*^*fl/fl*^ n=6, *AggrecanCreER*^*T2*^*;Ift88*^*fl/fl*^ n=8; 22 weeks, *Ift88*^*fl/fl*^ n=10, *AggrecanCreER*^*T2*^*;Ift88*^*fl/fl*^ n=7; 34 weeks, *Ift88*^*fl/fl*^ n=19, *AggrecanCreER*^*T2*^*;Ift88*^*fl/fl*^ n=22.

**S4.**
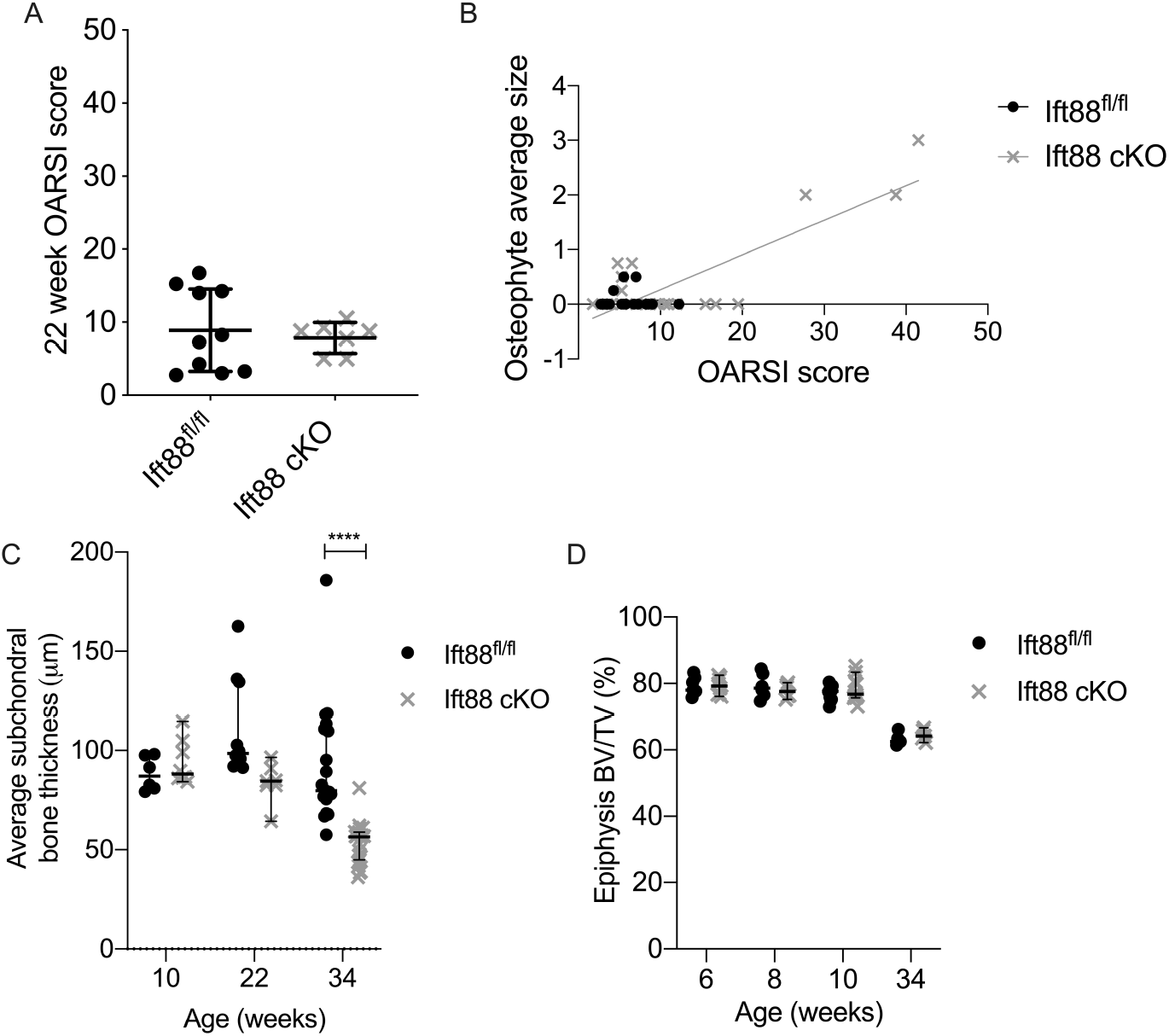
**A**, Summed cartilage OARSI scores in sections from 22 week old animals. *Ift88*^*fl/fl*^ n=10, *AggrecanCreER*^*T2*^*;Ift88*^*fl/fl*^ n=7. **B**, Osteophyte average size (data from Fig 4D) correlated with OARSI score in *Ift88*^*fl/fl*^ and *AggrecanCreER*^*T2*^*;Ift88*^*fl/fl*^ animals using simple linear regression *Ift88*^*fl/fl*^ n=10, *AggrecanCreER*^*T2*^*;Ift88*^*fl/fl*^ n=22. **C**, Median (average) subchondral bone thickness. 10 weeks, *Ift88*^*fl/fl*^ n=6, *AggrecanCreER*^*T2*^*;Ift88*^*fl/fl*^ n=8; 22 weeks, *Ift88*^*fl/fl*^ n=10, *AggrecanCreER*^*T2*^*;Ift88*^*fl/fl*^ n=7; 34 weeks, *Ift88*^*fl/fl*^ n=19, *AggrecanCreER*^*T2*^*;Ift88*^*fl/fl*^ n=22. **D**, Bone Volume/Total Volume (BV/TV) of epiphysis in *Ift88*^*fl/fl*^ and *AggrecanCreER*^*T2*^*;Ift88*^*fl/fl*^ animals between 6 and 34 weeks of age, n=5-10 in any group. Points represent median +/- 95% confidence intervals for each animal. Analysed by two-way ANOVA.

**S5.**
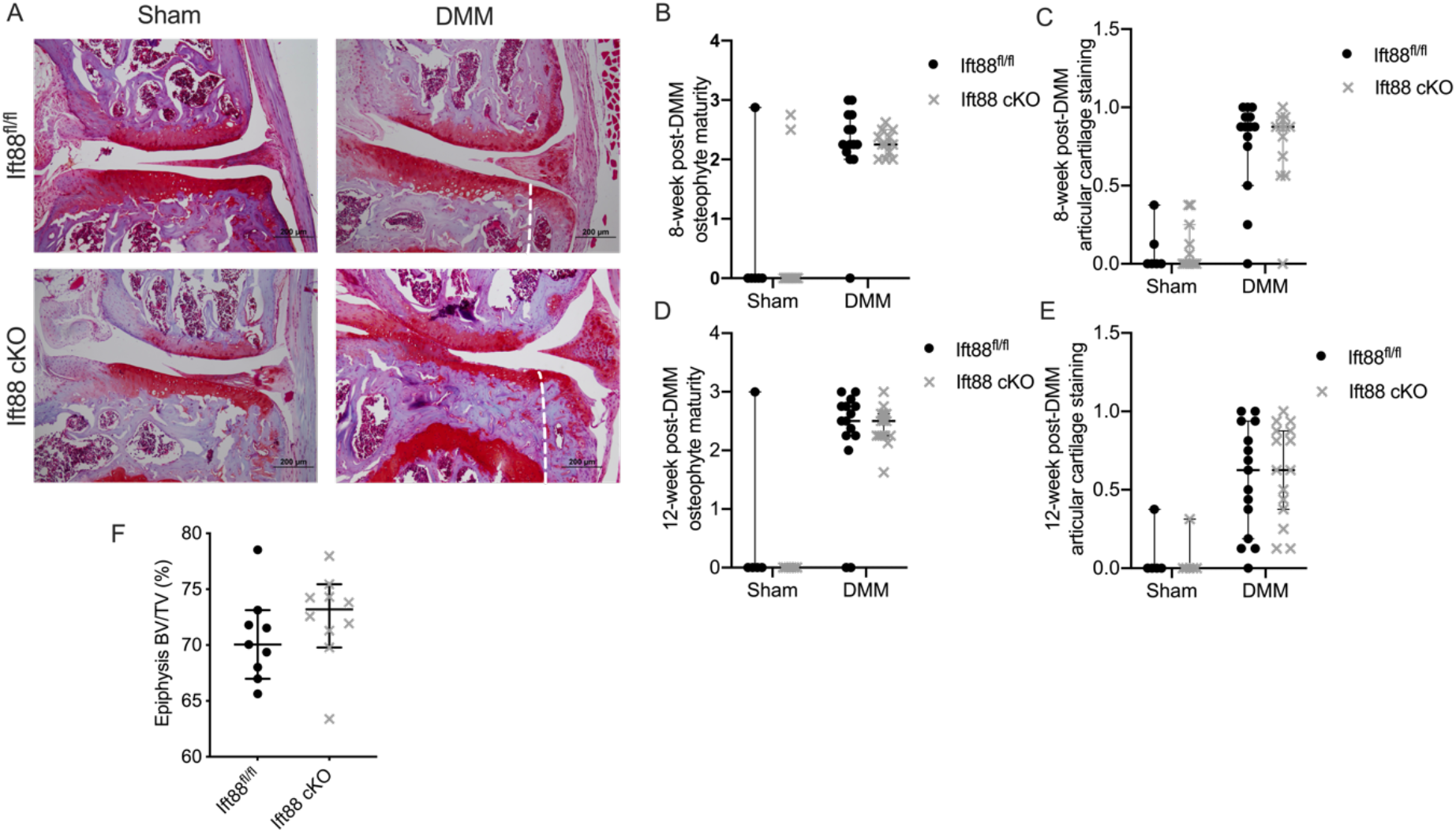
**A**, Safranin O stained histological sections of medial compartments from sham and DMM animals operated animals 12 weeks post-DMM surgery, (scale bar=200 μm). White dashed lines demarcate where osteophytes have formed. **B**, Osteophyte maturity score 8 weeks post-DMM. **C**, Articular cartilage staining score 8 weeks post-DMM. **D**, Osteophyte maturity score 12 weeks post-DMM. **E**, Articular cartilage staining score 12 weeks post-DMM. **F**, Bone volume/Total volume (BV/TV) of the epiphysis 12 weeks post-DMM in *Ift88*^*fl/fl*^ and *AggrecanCreER*^*T2*^*;Ift88*^*fl/fl*^ animals. 8 weeks post-DMM; *Ift88*^*fl/fl*^ n=14, *AggrecanCreER*^*T2*^*;Ift88*^*fl/fl*^ n=12. 12 weeks post-DMM; *Ift88*^*fl/fl*^ n=15, *AggrecanCreER*^*T2*^*;Ift88*^*fl/fl*^=15. Points represent median +/- 95% confidence intervals. Analysed by two-way ANOVA.

**S6.**
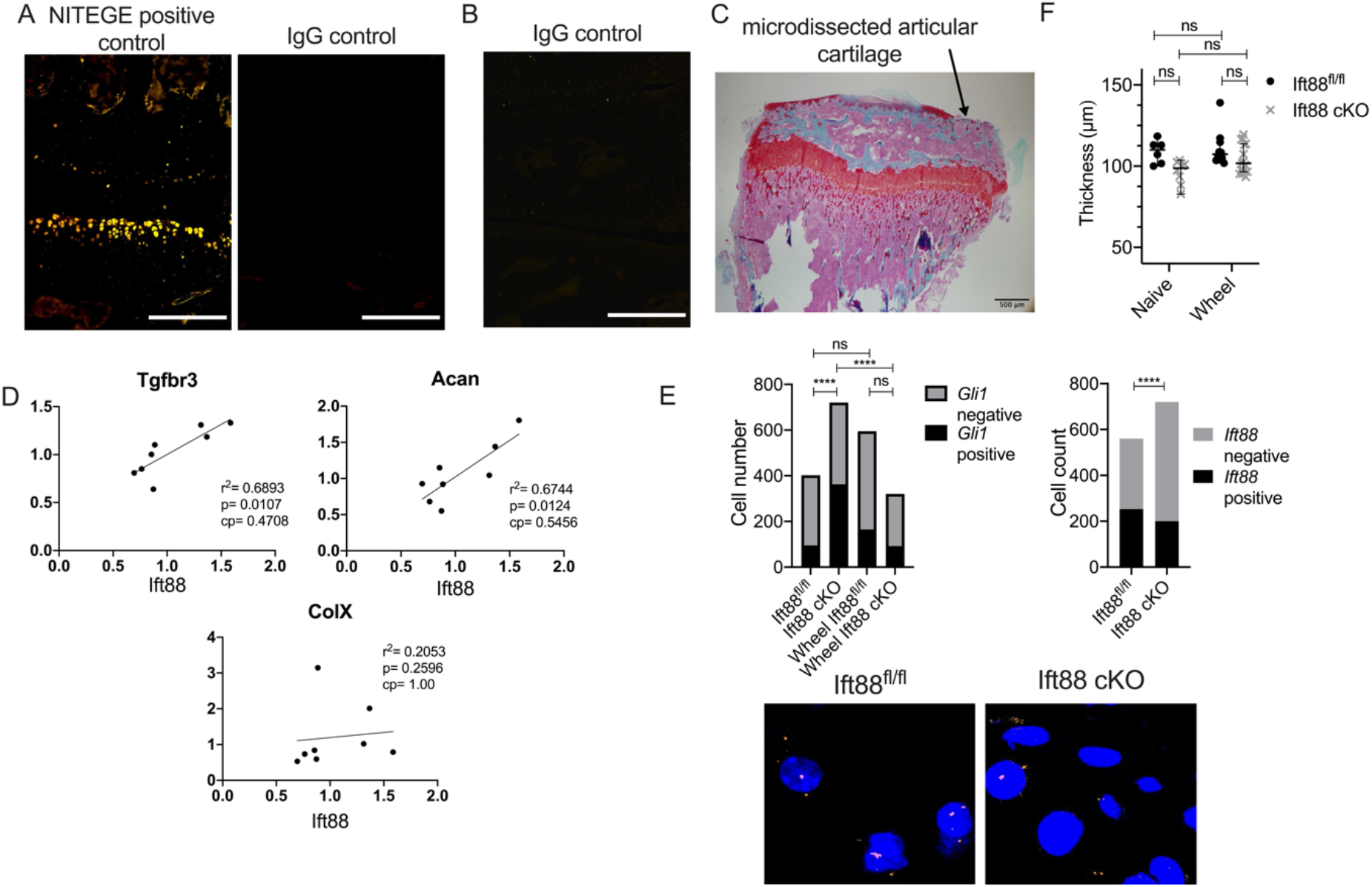
**A**, Medial compartment from animal 12 weeks post-DMM surgery was used to validate the NITEGE antibody (NITEGE positive control), and compared with IgG polyclonal. Scale bar=200μm **B**, Control IgG section for ColX staining from 10 week old animals (scale bar=200μm). **C**, Saf O staining of knee following micro-dissection of articular cartilage (scale bar= 500μm). **D**, RNA extracted from micro-dissected articular cartilage was analysed by qPCR to identify genes correlating with *Ift88* expression following normalisation to *Gapdh* and *Hprt*. Linear regression was performed and statistical significance assessed following Bonferroni correction (cp=corrected p value). *Tgfbr3* and *Acan* positively correlated with *Ift88* expression after correction whilst *ColX* did not correlate (see figure for p values and Supplementary Table 1). **E**, Contingency data of *Ift88* and *Gli1* positive nuclei in naïve *Ift88*^*fl/fl*^ and *AggrecanCreER*^*T2*^*;Ift88*^*fl/fl*^ animals and *Gli1* positive nuclei in wheel exercised (Analysed by Fisher’s exact test, ****p<0.0001). Representative images of *Ift88* positive nuclei. **F**, Cartilage thickness from naïve animals and mice that had access to wheel exercise (*Ift88*^*fl/fl*^ n=10, *AggrecanCreER*^*T2*^*;Ift88*^*fl/fl*^ n=15). Points represent median +/- 95% confidence intervals. Analysed by two-way ANOVA unless otherwise stated.

**Supplementary Table 1:**
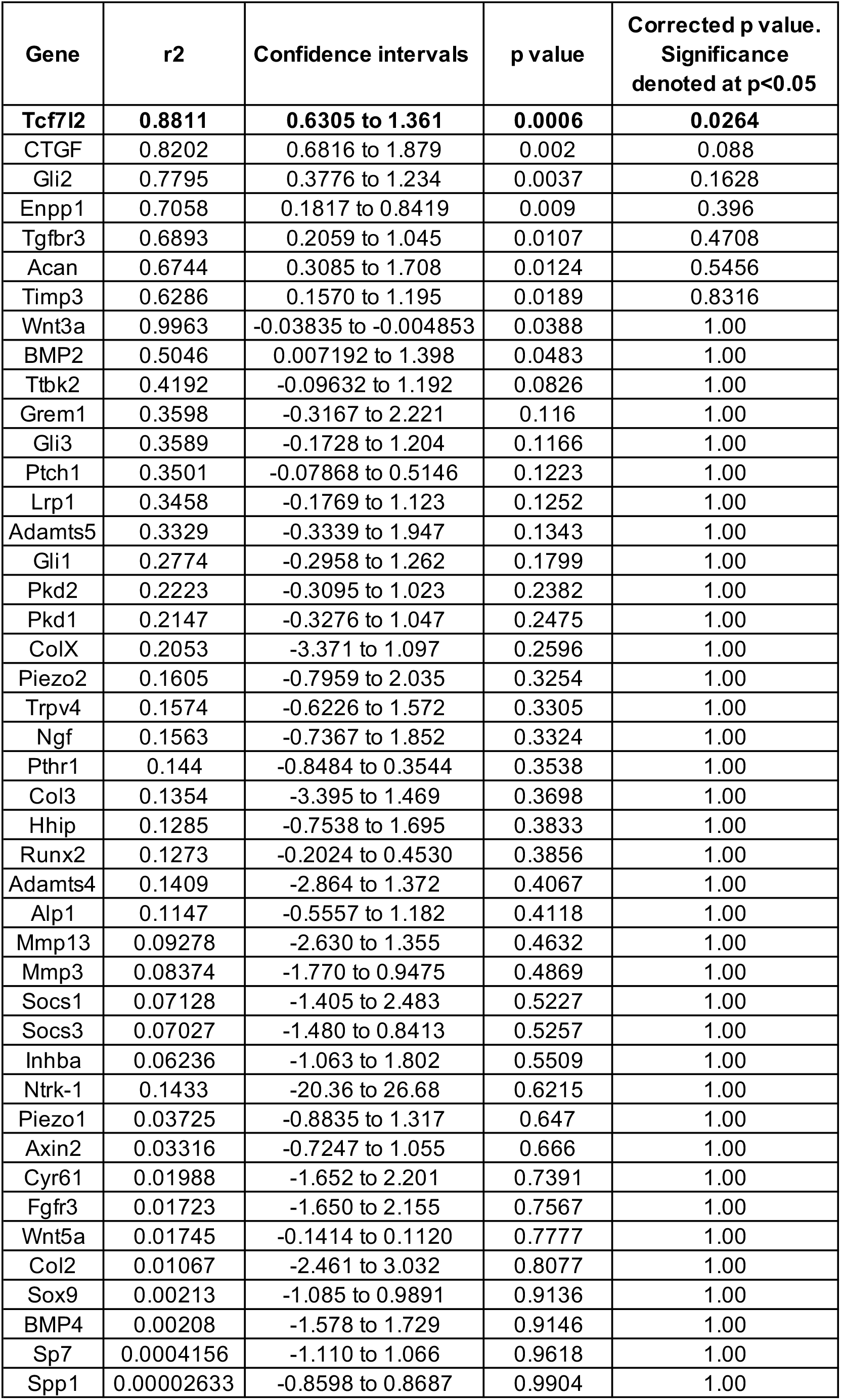
Average housekeepers of GAPDH and HPRT.

